## Supplementary Material for "DAT Val559 Mice Exhibit Compulsive Behavior Under Devalued Reward Conditions Accompanied by Cellular and Pharmacological Changes"

**Data Supplement:**

**Compulsive Behavior Under Devalued Reward Conditions Ac-companied by Alterations in Cellular and Dopamine Receptor Agonist Sensitivity in DAT Val559 Mice**

Adele Stewart PhD^1,2,#^, Gwynne L. Davis PhD^1,#^, Lorena B. Areal PhD^1^, Maximilian J. Rabil^1^, Vuong Tran^1^, Felix P. Mayer PhD^1^, and Randy D. Blakely PhD^1,2,*^

^1^Department of Biomedical Science and ^2^Brain Institute, Florida Atlantic University, Jupiter, FL.

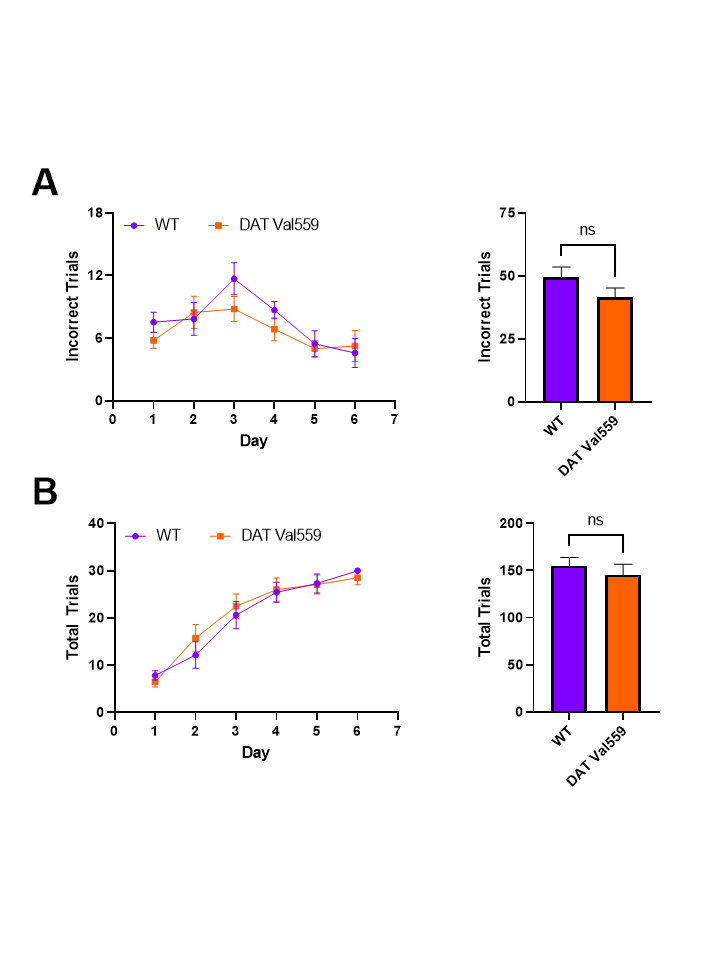
**Figure S1. Supplemental reversal learning data.** WT (n=14) and DAT Val559 (n=15) mice underwent pairwise discrimination training followed by a reversal phase. (A) Incorrect trials by day (left) and summed (right) during the reversal learning phase. (B) Trials completed by day (left) and summed (right) during the reversal learning phase. Data were analyzed by two-tailed student’s t-test or two-way RM-ANOVA with Sidak’s multiple comparison’s test. ns = not significant. Data are presented as mean ± SEM.

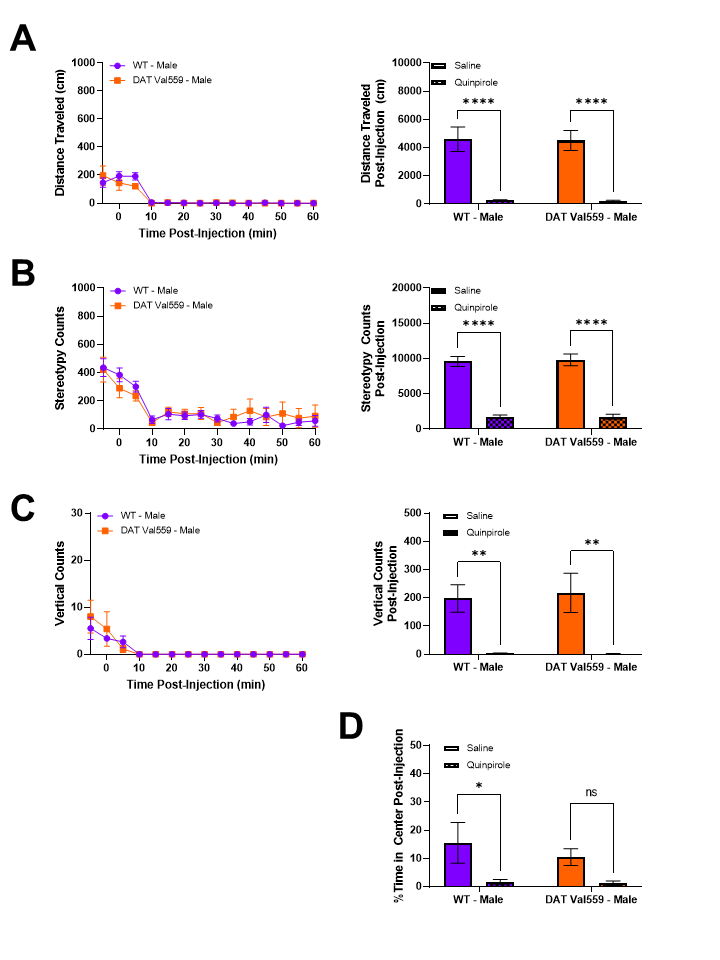

**Figure S2. Quinpirole-dependent locomotor suppression is comparable in WT and DAT Val559 mice.** WT (n=8) and DAT Val559 (n=8) mice were given a single injection of the D2/D3 agonist quinpirole (1 mg/kg, i.p.) and locomotor activity recorded for 60 minutes post-injection. Datasets are presented in 5-minute time bins across the recording period and as summary data adding up all activity post-injection. (A) Horizontal distance traveled. (B) Stereotypic motor movements. (C) Vertical locomotor activity. (D) % Time spent in the center of the chamber. Data were analyzed by two-way ANOVA with Sidak’s multiple comparison’s test. **P* < 0.05, ***P* < 0.01, ****P* < 0.001, *****P* < 0.0001. ns = not significant. Data are presented as mean ± SEM.

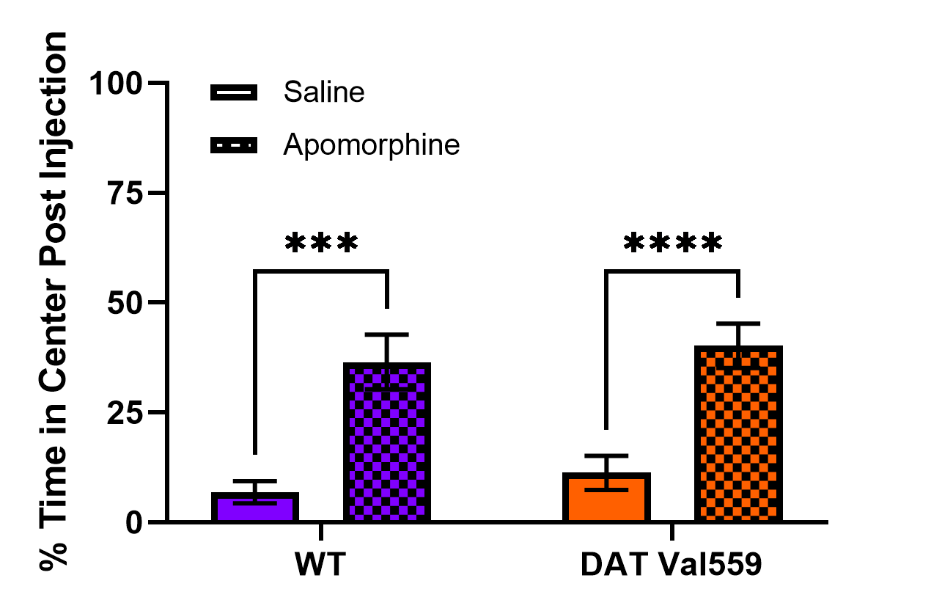

**Figure S3. WT and DAT Val559 mice exhibit an increase in center occupancy in response to apomorphine.** WT (n=13) and DAT Val559 (n=15) mice were given a single injection of the DA agonist apomorphine (5 mg/kg, s.c.) and locomotor activity recorded for 60 minutes post-injection. % Time spent in the center of the chamber after drug administration is depicted. Data were analyzed by two-way ANOVA with Sidak’s multiple comparison’s test. ***P < 0.001, ****P < 0.0001. Data are presented as mean ± SEM.

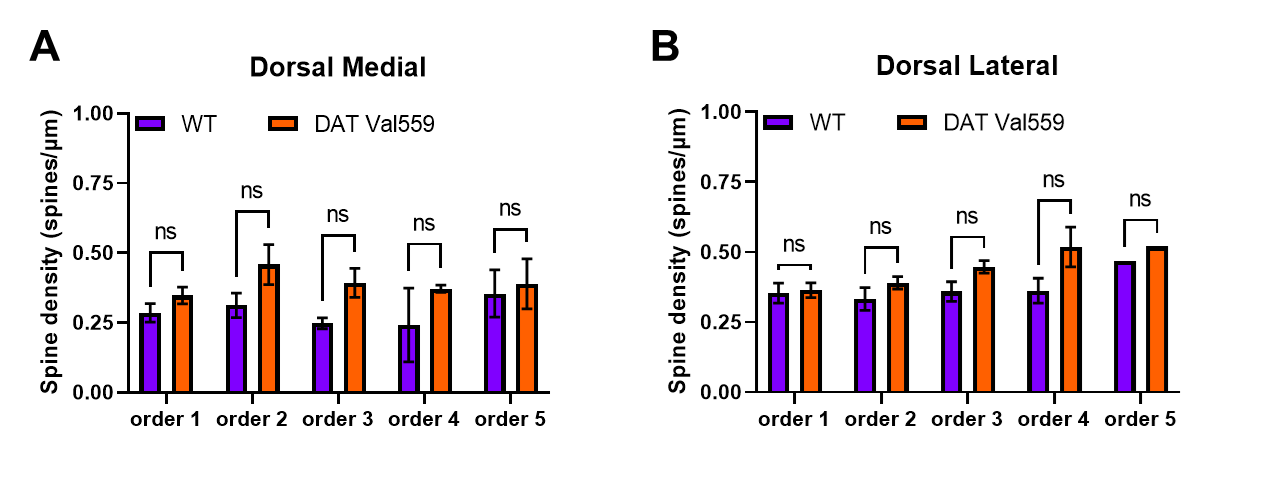

**Figure S4. Supplemental dendritic spine data.** Spine density in sections from WT (n=4) and DAT Val559 (n=4) mice was assessed utilizing Golgi staining coupled to brightfield microscopy. Spine densities were stratified based on degree of separation from the cell soma (order 1-5) in the (A) DMS and (B) DLS. Data were analyzed by two-way ANOVA with Sidak’s multiple comparison’s test. ns = not significant. Data are presented as mean ± SEM.
